# Local adaptation primes cold-edge populations for range expansion but not warming-induced range shifts

**DOI:** 10.1101/259879

**Authors:** Anna L. Hargreaves, Christopher G. Eckert

**Affiliations:** Department of Biology, Queen’s University, Kingston Ontario, Canada, K7L 3N6; Current address: Department of Biology, McGill University, Montreal Quebec, Canada, H3G 0B1

**Keywords:** range limits, experimental warming, phenology, local adaptation, reciprocal transplants, common garden

## Abstract

According to theory, edge populations may be the best suited to initiate range expansions and climate-driven range shifts if they are locally adapted to extreme edge conditions, or the worst suited to colonize beyond-range habitat if their offspring are genetically and competitively inferior. We tested these contrasting predictions by comparing fitness of low, mid, and high-elevation (edge) populations of the annual *Rhinanthus minor*, transplanted throughout and above its elevational distribution under natural and experimentally-warmed conditions. Seed from low-quality edge habitat had inferior emergence across sites, but high-elevation seeds were also locally adapted. High-elevation plants initiated flowering earlier than plants from lower populations, required less heat accumulation to mature seed, and so achieved higher lifetime fitness at high elevations. Fitness was strongly reduced above the range, but adaptive phenology enhanced the relative fitness of high-elevation seeds. Experimental warming improved fitness above the range, confirming climate’s importance in limiting *R. minor*’s distribution, but eliminated the advantage of local cold-edge populations. These results provide experimental support for recent models in which cold-adapted edge populations do not always facilitate warming-induced range shifts. The highest fitness above the range was achieved by a ‘super edge phenotype’ from a neighboring mountain, suggesting key adaptations exist at the regional scale even if absent from local edge populations. Our results demonstrate that assessing the value of edge populations will not be straightforward, but suggest that a regional approach to their conservation, potentially enhancing gene flow among them, might maximize species’ ability to respond to global change.

**Significance:** Individuals from range-edge populations are the most likely to disperse to habitat beyond the species current range, but are they best suited to colonize it? Our multi-year transplant experiment throughout and above the elevational range of an annual herb in the Canadian Rocky Mountains found that adaptive flowering phenology enhanced the fitness of high-edge seeds above the range, outweighing detrimental effects of poor seed quality. However, only one edge population maintained its advantage over central populations under experimental warming. While edge populations were most likely to drive range expansion, adaptation to cold climates may not help them initiate range shifts in response to climate warming, unless superior genotypes spread among isolated edge populations.

## Introduction

Decades of theory exploring the ecological and evolutionary process limiting species distributions yield contrasting predictions about range-edge populations. On one hand, species are thought to spread along environmental gradients via local adaptation at range margins (1). Along a continuous gradient, adaptation to the range edge should also confer an advantage beyond the range, priming edge populations to initiate future range expansions via niche expansion, and potentially facilitating range shifts in response to climate change (2, 3). However, theoretical models explaining stable range limits propose that environmental gradients reduce individual fitness and population size toward the range edge (4), an assumption increasingly supported by demographic surveys, transplant experiments and species distribution models (5-7). Under this scenario, offspring from small, isolated edge populations in harsh environments could suffer from the negative genetic effects of genetic drift and inbreeding, plus transgenerational environmental effects such as poor maternal provisioning. Low quality of offspring produced in edge populations could thwart adaptation, reduce their colonization ability, and reinforce range limits (5, 8). Thus, while individuals from the range edge are most likely to disperse beyond the range, whether they are the best or worst suited to colonize such habitat is unclear.

The potential for local adaptation in edge populations to promote range expansion along a stable environmental gradient is well established theoretically (9), with convincing examples from expansions of invasive plants (10). Whether local adaptation of edge populations will facilitate range shifts under climate change is less clear (11, 12). Cold-edge populations should initiate range shifts when their offspring have the highest fitness beyond the range. Edge propagules could gain such a fitness advantage from prior adaptation to a non-shifting gradient, e.g. photoperiod, or from adaptation to a shifting climate gradient if dispersal keeps pace with climate change such that beyond-range conditions always resemble the range edge more than the range center. If warming outpaces dispersal, however, cool-adapted edge populations may be less suited to colonize newly-warmed habitat than central genotypes, potentially stalling range shifts (13). Adaptation to cold climates can further undermine colonization ability if it involves a trade-off with potential fecundity, as commonly seen in plants (14). Cold-adapted plants often reproduce earlier but at a smaller size, increasing their absolute fitness in short growing seasons but reducing their relative fitness in longer ones (10).

Transplant experiments in natural environments are the best test of potential performance beyond a species range, local adaptation, and relative offspring quality (5). More than 50 years of over-the-edge transplant experiments show that many range limits are associated with declining habitat quality (5, 6), and reciprocal transplants commonly find local adaptation within species ranges (15). However, few reciprocal transplants are designed to test for adaptation towards and beyond range limits, and those that do include both central and edge source populations planted at the edge and beyond yield inconsistent evidence for local adaptation (16–18). Moreover, many transplants omit the early life history stages most closely related to offspring quality (19), and most do not replicate in time (multiple generations of lifetime fitness) or space (multiple edge and beyond-range sites), making results vulnerable to idiosyncratic site or year effects (5). Critically, none have included the relevant climate manipulations needed to reveal the role edge populations might play in responding to climate change.

We combined multi-year transplants with experimental warming to test the relative ability of edge populations of the annual herb, *Rhinanthus minor*, to initiate range expansion under natural conditions, and range shifts under climate warming. Along two transects spanning >1000 m of elevation (Nakiska ‘NK’, and Hailstone Butte ‘HB’), we reciprocally transplanted wild seed among populations well below the elevational range-center (‘low’), at the range center (‘mid’), and within 100 m elevation of the upper range edge (‘high’), and transplanted seed from these populations and two high-elevation populations on neighboring mountains to sites at and above the absolute range edge (‘edge’ and ‘above’; Fig. 1). Transplants were monitored from emergence to seed maturation to assess source differences in phenology and fitness. We test for the environmental and associated fitness gradients predicted to underlie many range limits, predicting that: a) habitat quality, measured by the lifetime fitness of the local source, declines at the range edge (5), b) low-quality edge habitat produces low-quality seeds, reflected in poor emergence across sites (19), and c) lifetime fitness is too low to sustain populations beyond the range. Using plant-height temperature sensors and climate station data, we calculate the growing season length and heat accumulation (growing degree days; GDD) at each site, and the GDD required for each source population to produce seed. We test for adaptation to elevation, predicting that: d) plants from sites with fewer GDD require fewer GDD to produce seed; e) at high elevations, edge populations achieve better post-emergence performance than other sources, and f) if adaptive traits that improve post-emergence performance outweigh low emergence, high-elevation sources will have the highest lifetime fitness at and above the range edge, priming them for range expansion. If local adaptation to cold environments involves the classic trade-off between early seed production and potential fecundity, high-elevation plants will reproduce earlier than other sources and be most successful at maturing seed, but produce relatively few seeds per plant. Finally, we subjected mid- and high-elevation seeds to experimental warming at and above the range edge to test whether g) temperature imposes the upper range limit, and h) edge seeds are best suited to colonize beyond-range habitat under warming.

**Fig. 1.**
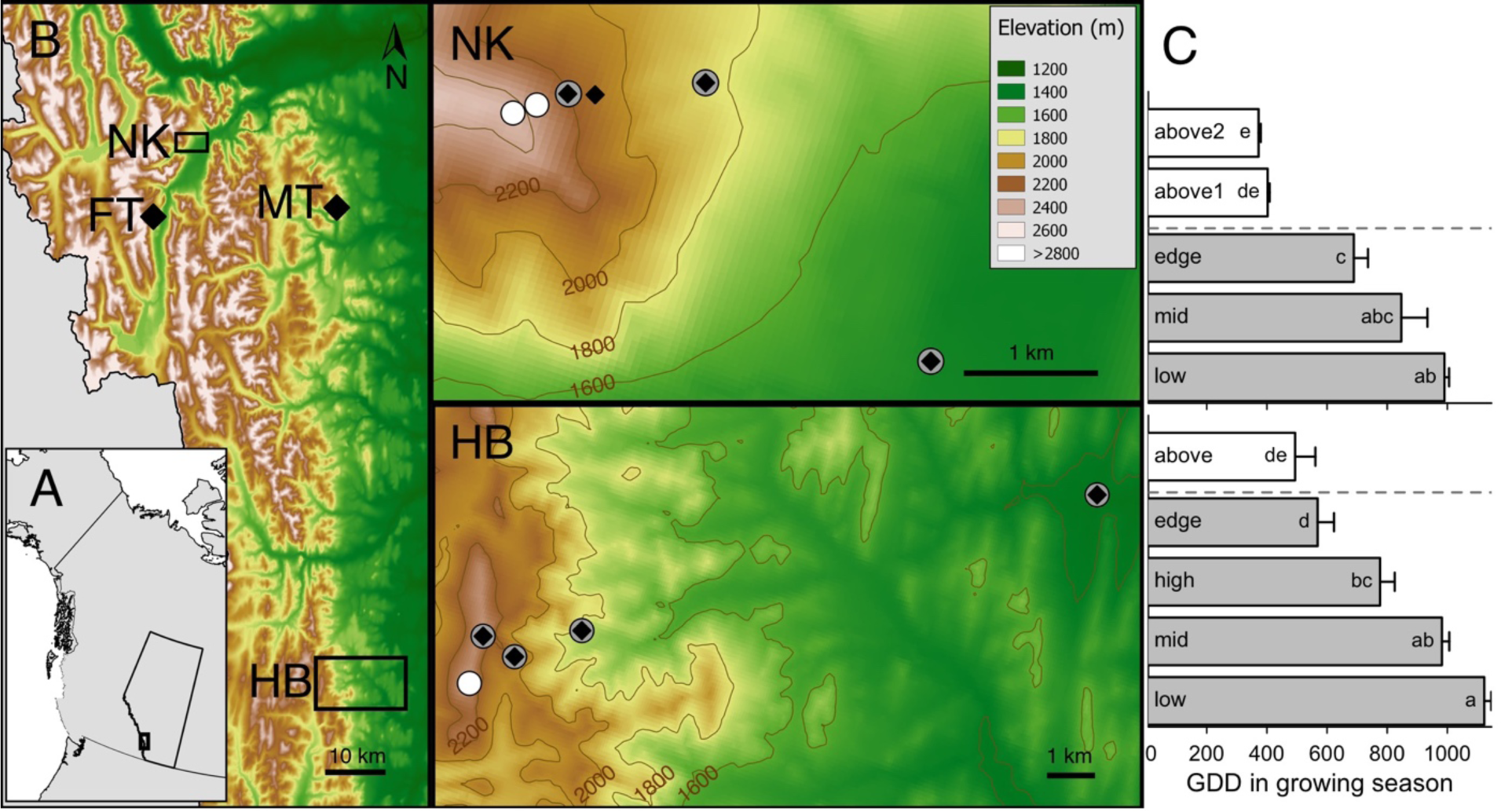
Location and climate of field sites. (*A*) Thick rectangle shows study area within Alberta, Canada, enlarged in *B. (B)* Elevation maps locating the four mountains that provided source material (left), with enlargements (right) showing reciprocal transplant transects: source populations (black diamonds), transplant sites within *R. minor’s* range (grey circles), transplant sites above *R. minor’s* upper range limit (white circles). (C) Mean ± SE heat accumulation per growing season at transplant sites during the 2011-2013 growing seasons. Dashed line indicates the upper range limit, contrasting letters indicate significant differences among sites and are comparable across transects (negative binomial GLM, likelihood ratio test of null model with intercept only vs. model with ‘site’ as a main effect; site 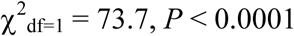).

## Results and Discussion

All measures of performance declined above *R. minor’s* upper range limit along both transects (Figs. 2&3, Table S1), even though fitness did not always decline from central to edge planting sites (Fig. 3, Table S2). Mean lifetime fitness (Fig. 3) was less than that required for self-sustaining populations at all sites above the range, and at the small outlier edge population that defined the absolute range limit on the HB transect. Even when seed dormancy was accounted for, low fitness translated to negative estimated growth rates (λ < 1) at the four highest sites (Fig. S1). The strong role of fitness constraints in imposing *R. minor’s* high-elevation range limit is consistent with most ecological and evolutionary models of stable range limits (4) and most empirical range limits studied to date (5, 6), and sets the stage for both local adaptation and poor quality of edge populations. As predicted, when edge habitat was of poorer quality than central habitat (i.e. lower fitness of local seeds at edge vs. central sites along HB; Fig. 3), edge seeds were also of lower quality, with significantly lower emergence across sites (HB; Fig. 2A).

**Fig. 2.**
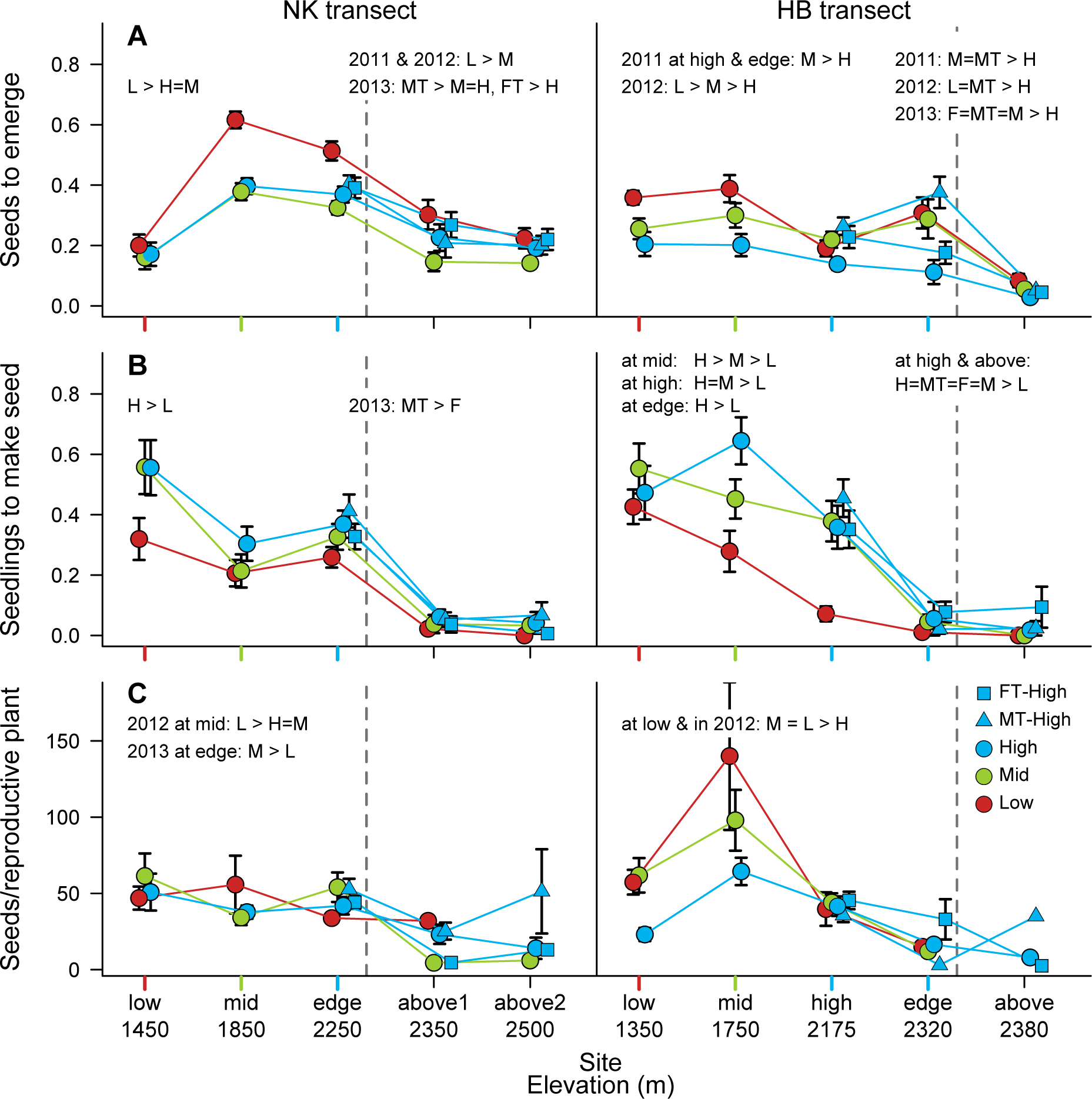
Performance of seed from different elevations within and above *R. minor’s* elevational range. Each performance parameter at each transect is analyzed as two experiments: 1) performance at sites within the range (left of dashed line) by local low-elevation (red circles), mid-elevation (green circles), and high-elevation (blue circles) sources, x-axis tick colour indicates the most local source population for each site; 2) performance across the high-elevation range limit (dashed line) including the highest three sites on each transect, comparing seeds from local sources and neighbouring high-elevation populations (blue squares and triangles). Text indicates significant differences among sources in the range (left justified) and across the range limit (spanning dashed line), separated by year and/or site if the final model contained significant source × year or source × site interactions. Full statistical results in Tables S1 and S2. Points are mean ± SE of performance across three yearly cohorts (2011, 2012 and 2013) for all sites except HB-edge (2011 only) and NK-above1 (2011 and 2012 only).

**Fig. 3.**
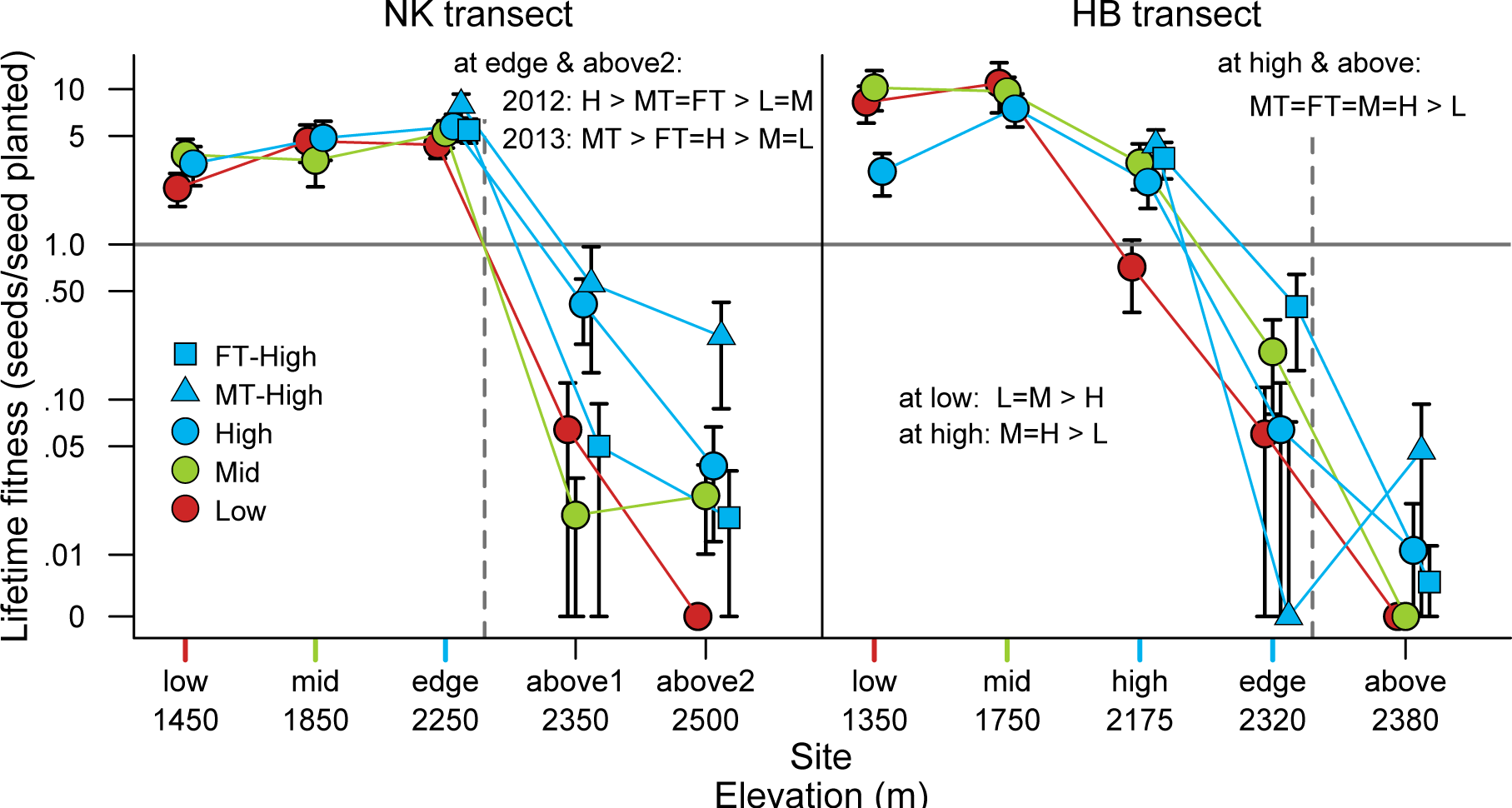
Mean ± SE annual lifetime fitness from 2011-2013 in and above *R minor’s* elevational range. Text shows significant differences among sources. At NK, source differences in performance at component life history stages (Fig. 2) did not result in local adaptation (i.e. a significant site × source interaction) within the range, but high-elevation sources (blue points) significantly outperformed Mid- and Low-elevation sources at and above the range limit. At HB, sources showed significant local adaptation within the range (site × source interaction 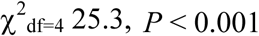, left-hand text), which was broadly reflected in performance across the upper range limit. Formatting as per Fig. 2, full statistical results in Tables SI and S2.

The sharpest decline in performance at sites too high to sustain populations was the proportion of emerged seedlings that matured seed (Fig. 2B). Indeed, all source populations failed to produce seed in at least one year at every above-range site (Fig. S2), which should impose severe viability constraints for annual species with only modest seed dormancy. Failure to mature seed was not due to poorer survival above the range, as plant longevity did not decline from high-elevations to above-range sites (Fig. S3). Rather, above the range many plants stayed small and never flowered, a phenomenon never observed within the range. Plants that did produce reproductive structures initiated them later than within the range (Fig. 4), often losing some or all to late-season damage from frost or snow. These results add to mounting evidence that reproductive failure sets many species’ range limits (5, 20).

**Fig. 4.**
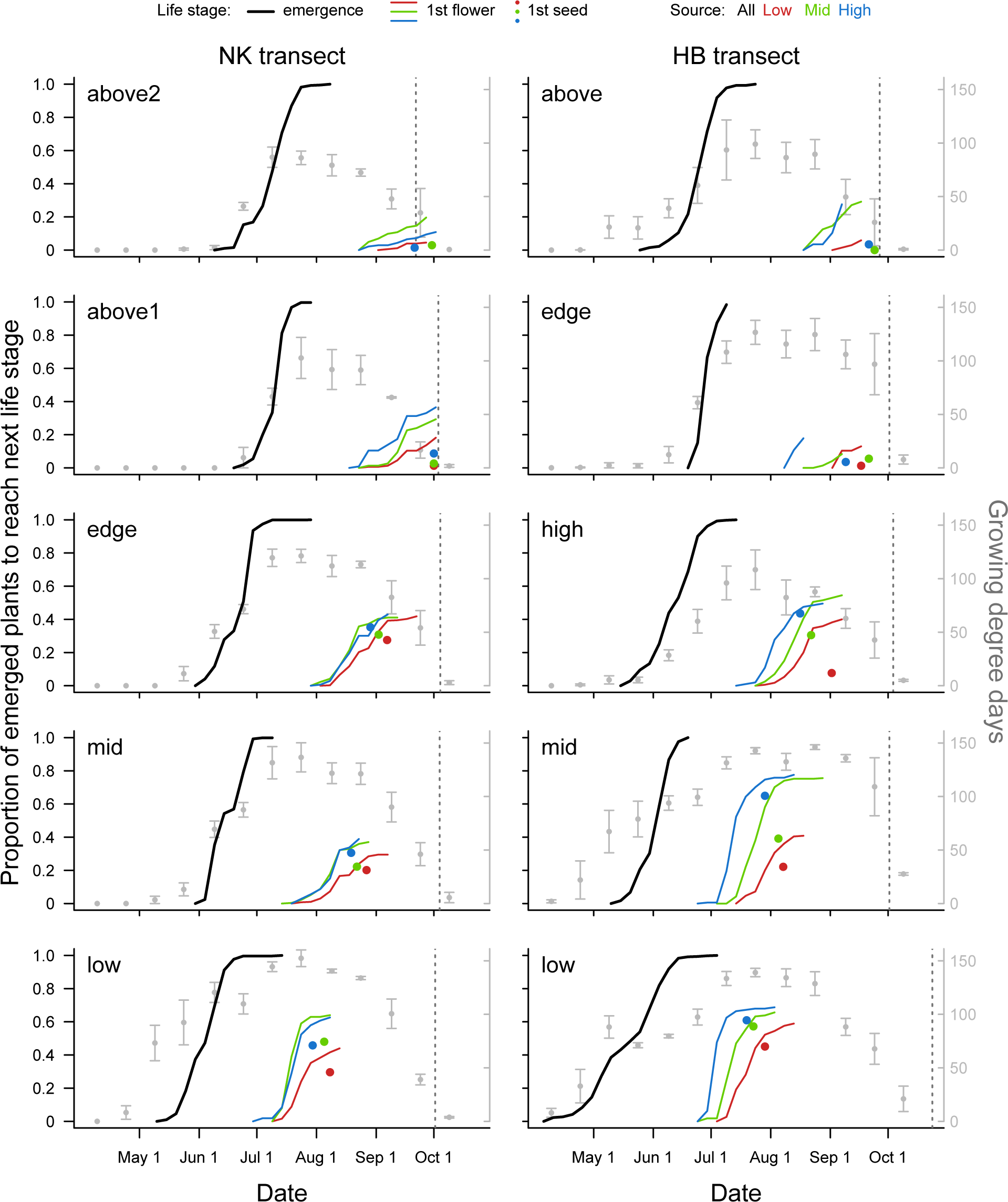
Phenology of plants from contrasting elevations transplanted across *R minor’s* range. Seeds were from low-, mid- and high-elevation populations local to each transect, transplanted to sites within (low to upper range edge) and above the elevational range. Curves show the accumulation of individual plants reaching a given life stage for 2011-2013 combined (2011-12 only for NK-above1, 2011 only for HB-edge), but models consider subplot means. Source populations did not differ in mean emergence date at any site so emergence curves combine data across sources (likelihood ratio tests, source contrasts *P >* 0.1, Table S3). Sources differed significantly in mean date of first flower both within and above the range. At NK plants from mid- and high-elevation seed flowered earlier than plants from low-elevation seed across sites (likelihood ratio tests, source *P* < 0.001 for each year). At HB, high-elevation plants flowered earlier than mid-elevation plants which flowered earlier than low-elevation plants across sites (likelihood ratio tests, source *P* < 0.001 for each year). Seed maturation was not monitored precisely enough to generate phenology curves, so colored points show the average date of first seed maturation across years for each source. Low points are missing from top panels as no low plants matured seed at the two highest sites. Seed maturation dates for NK-above1 are from 2012 only, as no seed was produced in 2011; all viable seeds were found during the last check in 2012 and differences in seed color (i.e. time since maturation) were not recorded, so sources have the same estimated seed maturation date. Grey points show heat accumulation (mean ± SE GDD for up to five years, 2010-2014) during each half of each month, dotted lines show mean estimated end of growing season defined by 3 days of consecutive snow pack or 2+ hours of −4° C or colder. Date of emergence and first flower differ among sites within transects, getting progressively later as elevation increases (Table S3).

Reduced ability to mature seed is consistent with the strong decline in growing season warmth, measured as growing degree days (GDD) per growing season, which decreased by 60% from the lowest to highest sites (Fig. 1C). Growing season GDD was the only climate variable that changed consistently across the range limit in the same manner as lifetime fitness (Fig. 1C; other climate variables shown in Fig. S4). The four sites above 2300 m, where fitness was below replacement, accumulated significantly fewer GDD than sites below 2300 m where transplanted populations were self-sustaining (Fig. 1C). The importance of growing season warmth in limiting above-range fitness is further corroborated by natural temperature variation among years. Compared to long-term averages, July and August 2012 and 2013 were unusually warm above the range at NK (Fig. S5) and seedlings from multiple sources matured seed, whereas none matured seed in the more typical summer of 2011 (Fig. S2). Indeed, in the warmest growing season of our study, 2013, there were 25% more GDD above the range at both transects compared to the coldest (2011; Table S3), and > 30 times more plants matured seed (9 of < 3000 seeds planted above the range in 2013 vs. 1 of > 7000 planted in 2011).

Predictable environmental gradients that limit fitness at and beyond range edges should impose directional selection on edge populations, resulting in differentiation of traits associated with the gradient (14). Consistent with this hypothesis and the importance of declining growing season warmth, plants from sites with the fewest GDD required the fewest GDD to produce seed wherever they were planted within the range (Fig. S6). High-elevation plants were the first to initiate flowering across sites (Fig. 4, Table S4), sometimes finishing flowering before low-elevation plants began. Their early start meant high-elevation plants were also the most likely to mature at least some seed across sites (Fig. 2B). In contrast, plants derived from low-elevation seed delayed flowering to grow secondary branches; this increased their potential number of fruits, but also increased the GDD required to produce seed (mean ± SE: NK-Low 590 ± 16, HB-Low 552 ± 20) compared to high-elevation plants from the same transect (NK-High 521 ± 20, HB-High 468 ± 18; effect of source: NK 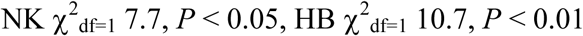, Fig. S6). Low-elevation plants thus produced more seeds given sufficient GDD (Fig. 2C), but high sources had greater lifetime fitness in the predictably shorter summers at high elevations (Fig. 3) and in growing seasons truncated by drought or cattle trampling at lower elevations (Fig. S7). Source differences in post-emergence phenology were enough to impart low- and high-elevation sources a home site-advantage consistent with local adaptation at one transect (HB).

Consistent with the hypothesis that local adaptation to a continuous environmental gradient primes edge populations for range expansion (5, 14), sources from high elevations, both local to each transect and from adjacent mountains, enjoyed the best performance above *R. minor’s* high range limit (Fig. 3). High-elevation plants outperformed low-elevation, and often mid-elevation plants in lifetime fitness (Fig. 3) and population growth rates derived from matrix models (Fig. S1) at all three above-range sites. Indeed, none of > 2000 low-elevation seeds planted ever made seed at the highest above-range sites (Fig. 2C). Thus, despite constraints on offspring quality, high-edge populations were best-suited to expand the species distribution under current conditions. The possibility that offspring from edge populations are simultaneously poor quality at early life stages and locally adapted later has been overlooked in the extensive theory on species distributions. In retrospect, their co-occurrence may be common since adaptation and offspring quality constraints arise from the same habitat and fitness gradients, and their interacting effects may result in counter-intuitive evolutionary dynamics at range edges (21).

Edge populations gained their above-range advantage from reproductive phenology, a consistently heritable functional trait (22) commonly involved in adaptation to climate (10), but which could also be influenced by non-heritable adaptive maternal effects. If adaptive maternal effects were solely responsible, the local advantage of high-elevation seeds would be short-lived - i.e. although seeds from low elevations had reduced relative and absolute fitness when planted at high elevations, their progeny might be as well suited to high-elevations as high-elevation genotypes. This would negate the evolutionary uniqueness of edge populations, eroding some of their conservation value. However, we feel that genetic adaptation to local environments is a more likely explanation. First, the most comprehensive review to date suggests adaptive maternal effects are generally uncommon and weak (23). Second, adaptive maternal effects are expected to evolve in response to conditions that are consistent at the seed dispersal (maternal) scale but variable at the scale of pollen dispersal (24). Conditions that are consistent at both scales, such as the large elevational differences studies here, should promote local adaptation instead (24). From an ecological perspective, the cause of edge-superiority is not particularly important: edge populations have value because they are the most likely and best able to colonize beyond-range habitat and expand the species range.

Experimental warming confirmed that inadequate heat accumulation at least partially limits *R. minor* fitness above its high-elevation range limit. Chambers warmed the air by 1.1 ± 0.18 °C (mean ± SD, Fig. S8; see SI for full discussion of OTC effects). This is roughly the temperature equivalent of descending to the next highest sites (Fig.s S4C and S8), so to the extent that elevational fitness patterns reflect temperature, fitness in OTCs should resemble fitness at the site below (Fig. 3). In line with these predictions, artificial warming increased performance at the highest above-range sites, but not enough to make populations viable (as at the second-highest sites: Figs. 5 and S1), and improved fitness at the HB-high but not the NK-edge site (Fig. 5). These results, combined with higher above-range fitness in naturally warmer years and the tidy relation between the GDD a source needs to set seed and the GDD available at its home site, clearly illustrate that growing season heat accumulation is important in setting *R. minor’s* high-elevation range limit. This does not preclude other features of the elevational gradient contributing to fitness patterns, though earlier experimental work rules out pollination deficits (25) and herbivory (26). We are often asked whether host plant abundance might limit *R. minor’s* distribution, just as the mycorrhizal community can influence range limits of non-parasitic plants (27). Although host genera are found above *R. minor’s* range (and transplants were always planted in patches of suitable hosts), the community does change across the upper range edge, including a decline in the relative abundance of legumes, which are thought to be particularly valuable (28). However, *R. minor’s* fitness did not covary with legume abundance at the local scale within or above the range (28).

**Fig. 5.**
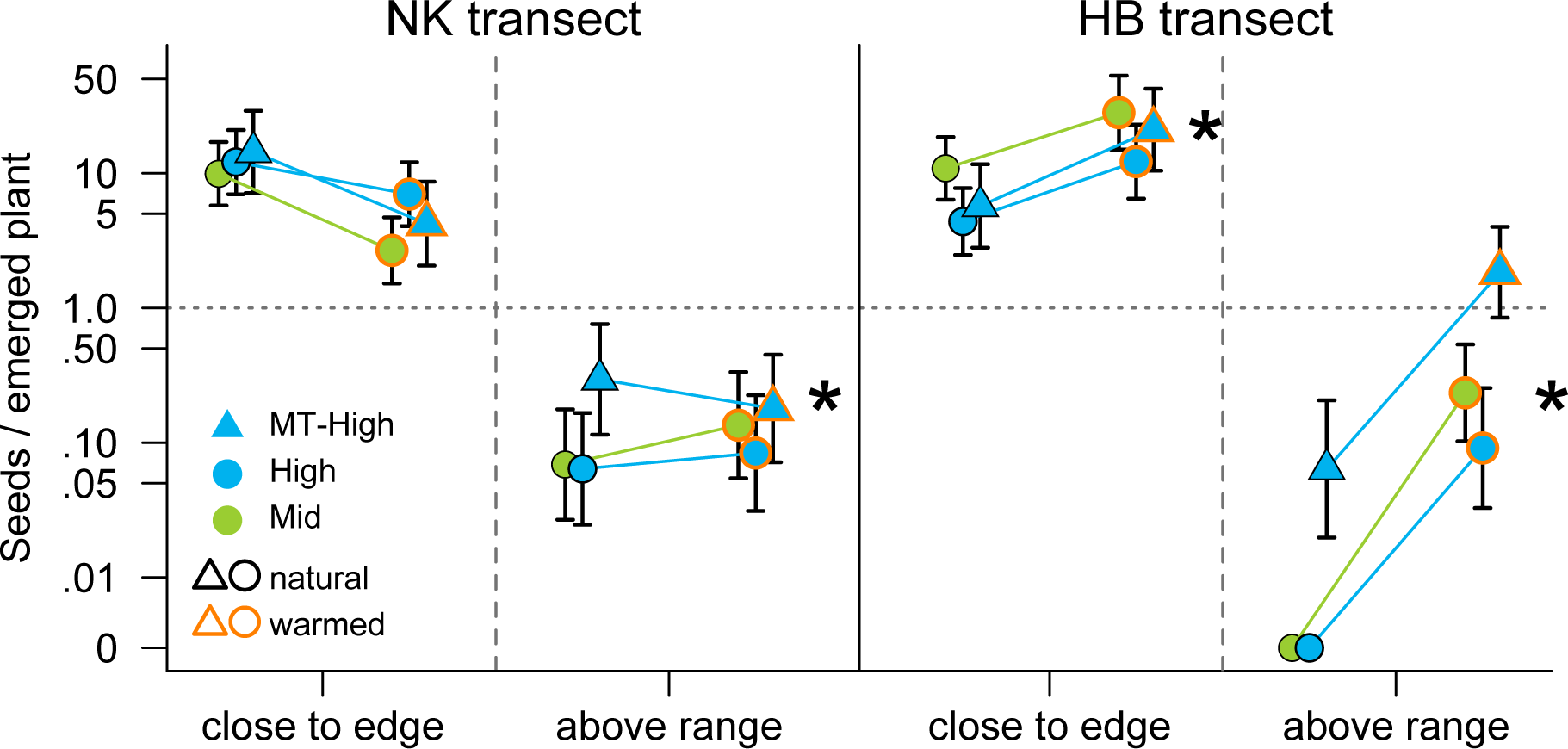
Effect of experimental warming. Plants in warmed plots outperformed those growing in natural climate conditions above the range at both transects, and close to the range edge at HB. The effect of treatment was significant but varied among sites and sources (minimum adequate model: treatment × site_type + treatment × transect + site_type × source, where ‘site_type’ was ‘close to edge’ or ‘above range’). Sites were NK-edge (2250 m), NK-above2 (2500 m), HB-high (2175 m) and HB-above (2380 m); vertical dashed lines denote upper range limit. * denotes sites where the effect of warming was significant (treatment × site interaction: likelihood ratio test, 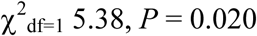). Overall the effect of warming was significant at the HB-transect but not the NK-transect (treatment × transect: 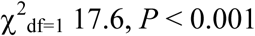), and seeds from the high population on neighboring Moose Mt outperformed local high and mid elevation seeds above the range (site × source: 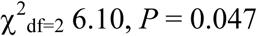). Points show least squared means ± SE back-transformed from the full model (site × source × transect × treatment + (1|year) + (1|subplotID), and combine data from 2012 and 2013.

Experimental warming also revealed the inconsistent effect of local adaptation. Although high sources outperformed low and mid-elevation sources at all three above-range sites (Figs. 3 & S1), local high seeds lost their advantage under experimental warming. When above-range plots were warmed, mid seeds had numerically (though not statistically) higher post-emergence performance (Fig. 5) and estimated population growth rates (Fig. S1) than plants from local high-elevation seeds. Mean temperatures and fitness in warmed above-range plots were still lower than at the range edge (Fig.s S8 & 5), so the advantage of central seeds was not because OTCs mimicked conditions in the range center. Local high seeds also failed to maintain their advantage over central seeds in the unwarmed control treatment (Fig. 5), which at first seems at odds with their overall superiority at high elevations (Fig. 3). This discrepancy arises because the warming experiment control treatment does not include data from 2011, when local high seeds had a particular advantage over mid seeds at HB (possibly because 2011 was colder than 2012-2013), or from the HB-edge and NK-above1 sites, where local high seeds again had a particularly strong advantage (perhaps because performance was just high enough for the advantage to be expressed in lifetime fitness, compared to the extremely low fitness at the highest sites; Fig. S2). The advantage of local edge populations was thus ephemeral, manifesting primarily in extreme environments and disappearing with even slight amelioration of the fitness-limiting gradient. Our empirical results support simulation model demonstrations that local range-edge populations may not be essential participants in warming-induced range shifts (13).

Warming chambers directly test whether cold temperatures constrain post-emergence fitness, but only partially mimic the effects of climate change — would other aspects alter our conclusions? We applied OTCs just as seedlings were emerging, whereas climate change will also advance spring melt (29) thereby adding growing season GDD and accentuating the effects of warming we observed. Climate change can also increase or decrease snow depth (29), which could improve or worsen above-range performance, respectively, if poor emergence (Fig. 2A) resulted from poor snow coverage at alpine sites (Fig. S4B). However, because there was no site × source interaction for emergence success (Tables S1&2) or timing (Table S4) in or above the range, climate change effects on emergence are unlikely to alter the relative success of seed sources. We did not measure OTC effects on summer soil moisture, but they tend to be neutral or slightly negative (30). Reduced soil moisture would likely have dampened the positive effect of warming, as *R. minor* is adversely affected by drought (Fig. S7). This may or may not emulate future climate change, which is predicted to increase precipitation but also evaporation in the Canadian Rocky Mountains (29, 31). Finally, under natural climate warming *R. minor* will not migrate upslope alone, and vegetation at our above-range sites may resemble vegetation in *R. minor’s* range more than it does currently. If anything, we expect this to have similar effects to warming, i.e. improving fitness for all source populations but reducing the advantage of high vs. mid-elevation seeds (Fig. 3). So, while OTCs do not perfectly mimic a climate-change scenario, we expect their main results — that warming improves fitness but eliminates the advantage of high seeds over mid-elevation seeds — to hold.

In contrast to the ephemeral advantage of local high-edge seeds, high-elevation seeds from neighboring Moose-mountain outperformed other sources in lifetime fitness (Fig. 3) and growth rates (Fig. S1) at five of the six highest sites, and in both natural and warmed plots of the OTC experiment (Fig. 5). Based on theory, the success of this population was unexpected; it was not larger, less isolated, different climactically, or more fecund at its home site than other high-elevation populations (Table S5; (32)). That this ‘super phenotype’ was restricted to one of four mountains, highlights that uniformly beneficial adaptations may be unable to spread among populations (33), particularly isolated high-elevation edge populations (16, 33). Genetic isolation of edge populations from each other, often detected in population genetics studies (34), likely plays an underappreciated role in stalling adaptation at range limits (35).

The conservation importance of range-edge populations is vigorously debated (3, 36), and our results add important insights to this broader conversation. Discouragingly, we show it may be difficult to assess the colonization potential, and thus the conservation value, of specific populations. Edge populations can harbor cryptic adaptations that facilitate success beyond the range even in the absence of a home-site advantage at the range edge (Fig. 3, NK transect). Superior populations may not be identifiable based on population size and isolation, parameters widely thought to determine population quality (37). Finally, although edge populations were best suited for natural range expansion, local-edge populations lost their advantage over central genotypes under climate warming, thus their importance for range dynamics may be context dependent. More hopefully, the global success of some non-local edge genotypes suggests that gene flow between isolated edge populations could enhance fitness both at and above the range edge (35). Considering these results together, we suggest that a regional approach to conserving isolated edge-populations, potentially including enhanced gene flow among them, could maximize species’ ability to respond to global change.

## Materials and Methods

### Study Species

*Rhinanthus minor* L. (Orobanchaceae) is native to Europe and North America, where it grows on various soils in open, often gently disturbed areas (38). It is a generalist root hemiparasite that obtains nutrients, but rarely photosynthate, from >50 host species, especially grasses and legumes (38, 39). Primary stems produce leaves in pairs. Each leaf node can produce either a secondary branch or a bud, which can develop into a single flower and fruit, thus there is a structural trade-off between growth and reproduction. Flowers are visited by bumble bees *(Bombus* spp.), but can produce a full complement of viable seeds – up to 18 per fruit – by autonomous self-pollination (25). Seeds rarely disperse > 2 m (38), making edge populations particularly critical for beyond-range colonization, though wet seeds can adhere to deer hide for >100 m, potentially enabling rare long-distance dispersal (Murphy L, Hargreaves AL, Eckert CG unpublished data). Most importantly for our purposes, *R. minor* is an annual with little seed dormancy, such that lifetime fitness can be measured every year across replicated range limits and is closely related to population demography. Within *R. minor’s* range 83 ± 7% (mean ± SD) of seeds emerged in their first spring, and dormancy did not differ among sources at sites where it could be monitored (high, edge and above-range sites; SI Materials and Methods), so we focus on fitness in the growing season after seeds were planted.

### Elevational Transects

We conducted reciprocal transplant experiments along two east-west elevational transects, 100 km apart (Fig. 1). Transects extended up consistent east-facing aspects of Nakiska (NK), and Hailstone butte (HB) peaks (Fig. 1, site details in Table S5). In our study area *R. minor* has an upper range limit around tree line (2200-2300 m), and a lower limit at ca. 1200 m. Potentially suitable habitat (open areas with grasses and legumes) occurred above *R. minor’s* range on both transects. Populations above 1400 m were in protected but mixed-use areas. Cattle grazed low- and mid-elevation populations along HB, where ranching has been ongoing since ~1900 (40). Grazing generally occurred after *R. minor* began setting seed, but cattle accidentally gained early access to HB-low in 2012 and HB-low and –mid in 2013, dramatically lowering fitness (Fig. S3). The structure of the range edge differed between transects. At NK the highest *R. minor* population (i.e. the upper range edge) still had ca. 1000 plants, whereas at HB the upper edge was set by a small outlier population of < 30 plants.

### Transplant Design

Along each transect we identified a source population from: below the elevational range center (Low); the elevational range center (Mid); and the upper range-edge (Edge; Fig. 1). Edge populations produced too few seeds to supply the 1500 seeds/yr needed for reciprocal transplants without potentially affecting demography. We planted Edge seeds into each range-edge site to properly assess home-source performance, but used seeds from the highest population of 2000+ plants (High; 100 m elevation below the absolute range edge on both transects) for reciprocal transplants. Edge and High seeds had near identical performance at edge sites (Fig. S9, Table S6), so we consider high-elevation seeds to accurately represent range-edge populations.

We established five transplant sites per transect. Within the range these were in the low, mid, and range-edge populations of each transect, and the high population at HB (Fig. 1; for clarity in figures and tables, population names are capitalized when referring to them as sources and lower case as sites). We reciprocally transplanted seeds among elevations within transects; seeds from NK low, mid and high populations were planted at NK sites, while HB seeds were planted along the HB transect. We established transplants above *R. minor’s* range in naturally open meadow at the highest elevation at that aspect on each mountain (HB-above and NK-above2), and an intermediate site between the mountain top and *R. minor’s* range edge at NK (NK-above1, Fig. 1). HB-above was only 60 m above *R. minor’s* upper range limit, so no intermediate site existed. Despite this small elevational difference, HB-above had an alpine climate and plant community like NK-above2, due to its exposed location on the butte top. HB-edge and NK-above1 were both sheltered by rock outcrops that reduce wind but trap snow, and so had warmer but sometimes shorter growing seasons that the top sites and grassier, subalpine vegetation (Fig. S5). To increase the genetic variation of seeds used to test the upper range limit, we collected seeds from two additional high-elevation (2000 m) populations on nearby mountains - Moose Mt (MT) and Fortress Mt (FT; Fig. 1) - and planted them at high, edge and above sites.

During late summer and early fall 2010-2012, we collected mature seed from 30-40 maternal plants per source population (10-15 plants at HB-edge). Plants were > 1 m apart to reduce relatedness. 10 filled seeds per plant were assigned to each transplant site until the donor had contributed to all sites or all seeds were used. For each transplant site, seeds were pooled across maternal families within source population. Seeds were planted < 40 d after collection into 5-20 plots/site/year (Table S7), situated in areas with grasses, sedges or legumes to provide suitable hosts even above the range (38). Before planting we removed naturally occurring *R. minor* within and around plots before they set seed. Each plot contained one subplot per source population, with 25 seeds planted 1 mm deep in a 5 × 5 cm grid (except 20 high edge seeds/subplot for HB-edge, as fewer seeds were available). Subplot order was randomized around the central stake that marked the plot. Subplots were separated by at ≥ 0.5 m except in warming chambers (see below) where they were closer due to space constraints. We marked the upper left corner of each subplot with a nail, and the outer seeds with toothpicks so seedlings could be readily located the following spring. Aside from removing naturally-occurring *R. minor*, the vegetation in each plot was left intact such that biotic or abiotic factors impinging on experimental plants were not altered in any way.

Transplant design varied slightly between years. For 2011, we planted 10 plots/site in 2010. For 2012, five extra plots were planted at HB-low to offset losses from cattle, but snow covered the HB-edge site before plots could be planted. For 2013, sample sizes were reduced to five plots at low and mid sites (only 1.5 of which were successfully planted at HB-low due to soil compaction by cattle), and HB-edge and NK-above1 were not replanted. For 2013, we also planted five plots of Edge seeds at NK-above2 to test whether seeds from the absolute range edge were more or less successful than seed from the highest populations of >2000 plants. There were insufficient HB-edge seeds to plant at HB-above. Sample sizes per source per site per year are fully described in Table S7.

### Transplant Monitoring

Plots were visited starting after snow melt (May – July) and once every 1-2 weeks thereafter until plants matured seed. Individual plants were identified according to their position on the planting grid, marked, and followed throughout their lives. Plants growing ≥ 2 cm ‘off-grid’ were considered potential contaminants and removed. For each subplot, we calculated the proportion of seeds to emerge, and the proportion of emerged seedlings to survive and produce viable seed. For each seed-producing plant we counted the viable fruits, the viable seeds per fruit in a representative sample of fruits (~25% of fruits/plant), final size (total leaf nodes), and estimated lifetime seed production (total fruits × mean viable seed/fruit). Seeds with a blackened center or exterior mold never germinated in greenhouse trials and were considered nonviable. Lifetime fitness was calculated per subplot as (total seeds produced)/(seeds planted).

We calculated several phenology parameters. Emergence date for each seedling was estimated as the date the seedling was first observed, less the number of primary stem nodes × the mean number of days required to grow a primary stem node. Mean growth rates were estimated at the plot-level, and so account for faster growth rates at warmer sites. Date of first flower was estimated as the date the first flower was observed minus 0.5 d for each previous flower or senesced flower (developing fruit) present. We chose 0.5 d as detailed observations (25) suggested 1 primary node (i.e. two flowers) open per day on average, and because it did not produce overestimated dates we knew to be too early from past visits. Analyses on the observed date of first flower yielded almost identical results (Table S4). Date of first seed maturation was estimated for each source at each site (monitoring was not precise enough to calculate it per plant) based on the first day seeds were counted or fruits were seen open and notes made on seed maturity.

### Experimental Warming

To test whether heat accumulation limits performance at and above the range limit, additional plots were planted for an experimental warming treatment in 2012 and 2013, at NK-edge, HB-high, NK-above2 and HB-above (Table S7). For 2012, we planted 10 plots of Mid and High seeds per site, but due to poor emergence of seeds from the HB-High population, we decided to keep seven of the ten OTC-intended plots in the control treatment at the HB-high site (i.e. only put OTCs on three plots). In 2013, we added a third source, MT-High, at all sites. We also planted an extra 10 plots of Mid and High seed at HB-above to ensure enough emerged plants for the experiment; 2012 warming chambers were placed only on plots where seedlings emerged. Once emergence had begun at a given site, experimental plots were enclosed in open-topped warming chambers (OTCs) to passively warm the air around transplants. OTCs were clear plastic cones, 0.4 m tall with a 1.2 m diameter base and 1 m diameter opening (Supplementary methods). Chambers warmed the air at plant height by 1.1 ± 0.18 °C (mean ± SD), adding 1.4 ± 0.43 GDD per day (Fig. S8).

### Climate Monitoring

We used HOBO (Prov v2; Onset) and iButton (DS1921G; Maxim) sensors to measure temperature at plant height (2 cm from the ground) at each transplant site (SI Materials and Methods). From these data, we estimated the days of insulating snowpack, minimum winter temperature, the start and end date of the growing season, and the growing degree days (GDD) with T_base_ of 10 °C and T_max_ of 30 °C in each growing season (calculation details in SI). Due to sensor destruction by lightning and animals we could not calculate all parameters in all years at all sites. Additional iButtons were deployed in 2012 and 2013 to compare temperature in control vs. warmed plots at the four sites in the warming experiment (*n* = iButtons per treatment across sites and years).

To compare climate among years we used publically available data from permanent weather stations. To assess differences in above-range climate during our study, we used air-temperature records from provincial or university-maintained weather stations at the same elevation and within 500 m of transplant plots at NK-above2 and HB-above. To assess whether temperatures encountered during our study were typical of long term climate, we used long-term records from the federal (Environment and Climate Change Canada) weather stations < 20 km from our sites with records including 2011-2013 (SI Materials and Methods).

### Analysis

Performance analyses consider subplot as the unit of replication, accounting for nonindependence of the 20-25 seeds within subplots. Emergence, proportion of emerged seedlings to reproduce, seeds produced/emerged seedling (post-emergence performance, warming experiment), and lifetime fitness (seeds produced/seeds planted) are calculated per subplot. Analyses of lifetime seed production (seeds produced/reproductive plant) and phenology use subplot means. Analyses of emergence combine data from control and warmed plots when both are available, as warming manipulations began after emergence.

We conducted three analyses of spatial patterns in performance: (i) patterns within the range, (ii) performance across the range limit, and (iii) the effect of experimental warming. All analyses were slightly complicated as site, source and year were not fully crossed. Within-range analyses were conducted separately by transect, and included home-transect sources (Low, Mid, High) at within-range sites that were planted in all three years (low, mid, edge for NK; low, mid, high for HB: full model `~ elevation × source × year`). We ran an additional model for HB comparing high and edge sites in 2011 (full model `~ elevation × source`). Analyses of performance across the range limit were conducted separately by transect, and included replicated home-transect sources (Low, Mid, High), foreign high-elevation sources (MT-High, FT-High), and sites spanning the upper range limit (edge, above1, above2 for NK; high, edge, above for HB) for three years (2011-13). NK-above1 and HB-edge sites were not planted every year, so we ran one model comparing all years excluding these sites (full model `~ elevation × source × year`), and one model including the intermediate sites for the years they were studied (NK full model `~ elevation × source × year` for 2011 and 2012; HB full model ~ elevation × source for 2011). Sources frequently failed to produce seeds above the range; accordingly we did not analyze seed production per reproductive plant.

We used generalized linear models (GLMs) with binomial error distributions for proportional parameters (logit link function; glm, MASS package(41)), negative binomial or Poisson distributions for parameters that include seed counts (log link; glm or glm.nb, MASS package), and linear models or Poisson GLMs for phenology of emergence and start of flowering, in R (3.3.3 2017-03-06 (42)). Initial models included all possible interactions among predictors (site, source, year). Significance of terms was assessed using type III sums of squares (Anova function, car package 2.1-4 (43)), except for negative binomial models as Anova does not recalculate the dispersion parameter; for these we compared models with and without a given term (likelihood ratio test using a χ^2^ distribution). Effects that were not intrinsic to the experimental test of our hypotheses (i.e. interactions and the effect of ‘year’), were dropped from models if not significant (P > 0.05). ‘Site’ and ‘source’ were integral to the experimental design and therefore retained in final models even if not significant. Another approach would have been to include ‘year’ and its interactions as random effects, but mixed models rarely converged. The validity of the final model was confirmed if it did not differ in explanatory power from the full model (likelihood ratio test, χ^2^ distribution). Binomial and Poisson GLMs assume a dispersion parameter (φ) of 1; φ was calculated for the minimum adequate model and those with f substantially different from 1 (<0.6 or >1.3) were re-run using a quasi distribution with further model simplification if warranted (44). Model selection using AICc or QAICc (dredge function, MuMIn package 1.15.6(45)) yielded similar results.

For significant non-interacting predictors, we tested for differences among factor levels using least squared means with the Tukey method to maintain an overall a = 0.05 (lsmeans function, lsmeans package 2.25 (46)). For main effects involved in significant interactions, pairwise tests were conducted within levels of the interacting term. If parameters were uniformly zero for a level of a main effect (i.e. total failure at sites above the range), significant pairwise differences between sites were identified when the back-transformed 95% confidence limits for non-zero-performance sites did not overlap with zero.

Warming experiment analyses consider the two transects together, with sites grouped as ‘close to the range edge’ (NK-edge, HB-high) or ‘above the range’ (NK-above2, HB-above). We used generalized linear mixed models that included a random intercept for year, and, to account for overdispersion, an observation-level random effect ‘subplotID’ (47); full model: `performance ~ treatment × source × transect × grouped_site + (l|year) + (1|suplotID)`. Interactions and ‘transect’ were dropped from models if deemed nonsignificant using likelihood ratio tests. As warming began after emergence, analyses compare post-emergence performance only (seeds / emerged seedling), which reflects maximum possible lifetime fitness (i.e. if all seeds emerged). Data include only sources involved in the experiment each year (local Mid and High sources for 2012, Mid, High and MT-High for 2013).

To assess whether sites spanning the range edge would sustain populations of each source given their overall performance and the observed level of seed dormancy, we calculated the density-independent population growth rates (λ) using a matrix model (48) (details in SI). Models used across-year means for emergence and seeds/emerged seedling for each source × site combination, and the overall transect mean for dormancy rates (i.e. across the three sites and all sources and years) as dormancy did not differ among sites or sources (SI). Finally, we estimated λ using post-emergence performance from the warming treatment and emergence data local to each site to test whether warmer growing seasons would make habitat suitable above the range. For above-range sites we also estimated λ using emergence data from the range edge to test whether warmer growing seasons combined with improved ameliorated emergence conditions would make habitat suitable.

Analyses of climate variables used negative binomial GLMs for site-level comparisons of GDD / growing season, and linear mixed models, with plot ID as a random intercept, for comparing temperature and GDD / day between control and warmed plots (Supplementary methods).

## Acknowledgements

We are grateful to numerous field assistants who helped with the 20 months of fieldwork, especially L. Falk, E. Bothwell, and G. Langston, to G. Langston again for help creating the map in Fig 1, and to B. Leung for help with matrix models. This research was funded by the Natural Sciences and Engineering Research Council (NSERC) of Canada Discovery Grant to C.G.E, an Alberta Conservation Association Biodiversity Grant to A.L.H, and NSERC CGS, Queen’s University and IODE Canada doctoral scholarships to A.L.H.

## Author contributions

The study was jointly designed by the authors. A.L.H conducted the field work and data analyses and drafted the manuscript, C.G.E contributed substantial editing to the final version.

## References

1. Levin DA (2000) The origin, demise, and expansion of plant species (Oxford University Press, New York).

2. Hunter ML & Hutchinson A (1994) The virtues and shortcomings of parochialism - Conserving species that are locally rare, but globally common. Conserv. Biol. 8(4): 1163–1165.

3. Gibson SY, Van der Marel RC, & Starzomski BM (2009) Climate change and conservation of leading-edge peripheral populations. Conserv. Biol. 23(6):1369–1373.

4. Sexton JP, McIntyre PJ, Angert AL, & Rice KJ (2009) Evolution and ecology of species range limits. Annu. Rev. Ecol. Evol. Syst. 40:415–436.

5. Hargreaves AL, Samis KE, & Eckert CG (2014) Are species’ range limits simply niche limits writ large? A review of transplant experiments beyond the range. Am. Nat. 183(2): 157–173.

6. Lee-Yaw JA, et al. (2016) A synthesis of transplant experiments and ecological niche models suggests that range limits are often niche limits. Ecol. Lett. 19(6):710–722.

7. Pironon S, et al. (2017) Geographic variation in genetic and demographic performance: new insights from an old biogeographical paradigm. Biol. Rev. 92(4):1877–1909.

8. Brown JH, Stevens G, & Kaufman DM (1996) The geographic range: size, shape, boundaries and internal structure. Annu. Rev. Ecol. Syst. 27:597–623.

9. Holt RD (2003) On the evolutionary ecology of species’ ranges. Evol. Ecol. Res. 5(2):159–178.

10. Colautti RI & Barrett SCH (2013) Rapid adaptation to climate facilitates range expansion of an invasive plant. Science 342(6156):364–366.

11. Hargreaves AL, Bailey SF, & Laird RA (2015) Fitness declines toward range limits and local adaptation to climate affect dispersal evolution during climate-induced range shifts. J. Evol. Biol. 28:1489–1501.

12. Bocedi G, et al. (2013) Effects of local adaptation and interspecific competition on species’ responses to climate change. Ann. N. Y. Acad. Sci. 1297:83–97.

13. Atkins KE & Travis JMJ (2010) Local adaptation and the evolution of species’ ranges under climate change. J. Theor. Biol. 266(3):449–457.

14. Hoffmann AA & Blows MW (1994) Species borders: ecological and evolutionary perspectives. TREE 9(6):223–227.

15. Hereford J (2009) A quantitative survey of local adaptation and fitness trade-offs. Am. Nat. 173(5):579–588.

16. Samis KE, Lopez-Villalobos A, & Eckert CG (2016) Strong genetic differentiation but not local adaptation toward the range limit of a coastal dune plant. Evolution 70(11):2520–2536.

17. Geber MA & Eckhart VM (2005) Experimental studies of adaptation in *Clarkia xantiana*. II. Fitness variation across a subspecies border. Evolution 59(3):521–531.

18. Stanton-Geddes J, Tiffin P, & Shaw RG (2012) Role of climate and competitors in limiting fitness across range edges of an annual plant. Ecology 93(7):1604–1613.

19. Donohue K (2009) Completing the cycle: maternal effects as the missing link in plant life histories. Phil. Trans. R. Soc. B 364(1520):1059–1074.

20. Griffith TM & Watson MA (2006) Is evolution necessary for range expansion? Manipulating reproductive timing of a weedy annual transplanted beyond its range. Am. Nat. 167(2): 153–164.

21. Gilbert KJ, et al. (2017) Local maladaptation reduces expansion load during range expansion. Am. Nat. 189(4):368–380.

22. Hendry AP & Day T (2005) Population structure attributable to reproductive time: isolation by time and adaptation by time. Mol. Ecol. 14(4):901–916.

23. Uller T, Nakagawa S, & English S (2013) Weak evidence for anticipatory parental effects in plants and animals. J. Evol. Biol. 26(10):2161–2170.

24. Galloway LF (2005) Maternal effects provide phenotypic adaptation to local environmental conditions. New Phytol. 166(1):93–99.

25. Hargreaves AL, Weiner JL, & Eckert CG (2015) High-elevation range limit of an annual herb is neither caused nor reinforced by declining pollinator service. J. Ecol. 103:572–584.

26. Falk L, Hargreaves AL, & Eckert CG (2013) The intensity and effects of insect leaf herbivory on *Rhinanthus minor* along its elevational range. BSc Honours (Queen’s University).

27. Afkhami ME, McIntyre PJ, & Strauss SY (2014) Mutualist-mediated effects on species’ range limits across large geographic scales. Ecol. Lett. 17(10): 1265–1273.

28. Bocchinfuso S, Ensing DJ, & Eckert CG (2017) Variation in beneficial host availability and performance of hemi-parasite *Rhinanthus minor* across and above its elevational range. BSc Honours thesis (Queen’s University, Kingston).

29. Barnett TP, Adam JC, & Lettenmaier DP (2005) Potential impacts of a warming climate on water availability in snow-dominated regions. Nature 438(7066):303.

30. Marion GM, et al. (1997) Open-top designs for manipulating field temperature in high-latitude ecosystems. Glob. Change Biol. 3:20–32.

31. Canadian Centre for Climate Modelling and Analysis (2017) Climate forecasts: Canada. (Environment and Climate Change Canada).

32. Hargreaves AL (2014) Evolutionary ecology of range limits: conceptual syntheses and empirical tests. PhD PhD (Queen’s University, Kingston, ON).

33. Leimu R & Fischer M (2008) A meta-analysis of local adaptation in plants. PLoS One 3(4): 1–8.

34. Eckert CG, Samis KE, & Lougheed S (2008) Genetic variation across species’ geographic ranges: the central-margin hypothesis and beyond. Mol. Ecol. 17:1170–1188.

35. Sexton JP, Strauss SY, & Rice KJ (2011) Gene flow increases fitness at the warm edge of a species’ range. PNAS 108(28):11704–11709.

36. Channell R & Lomolino MV (2000) Dynamic biogeography and conservation of endangered species. Nature 403(6765):84–86.

37. Hoffmann AA, Sgró CM, & Kristensen TN (2017) Revisiting adaptive potential, population size, and conservation. TREE 32(7):506–517.

38. Westbury DB (2004) *Rhinanthus minor* L. J. Ecol. 92(5):906–927.

39. Cameron DD, Coats A, & Seel W (2006) Differential resistance among host and non-host species underlies the variable success of the hemi-parasitic plant *Rhinanthus minor*. Ann. Bot. 98:1289–1299.

40. Tosh K (1975) Our Foothills (Millarville, Kew, Priddis and Bragg Creek Historical Society, Calgary).

41. McCullagh P & Nelder JA (1989) Generalized linear models (Chapman and Hall, London) 2nd Ed.

42. R Development Core Team (2015) R: a language and environment for statistical computing. (http://www.r-project.org/, Vienna, Austria).

43. Fox J & Weisberg S (2011) An R companion to applied regressionThousand Oaks CA: Sage).

44. Crawley M (2013) The R book (John Wiley & Sons, Chichester, UK) 2nd Ed.

45. Bartón K (2016) MuMIn: Multi-model inference. R package version 1.15.6.

46. Lenth RV (2016) Least-Squares Means: The R Package lsmeans. J. Stat. Softw. 69(1): 1–33.

47. Bolker BM (2015) Linear and generalized linear mixed models. Ecological statistics: Contemporary theory and application, eds Fox GA, Negrete-Yankelevich S, & Sosa VJ (Oxford University Press, Oxford), 1st Ed.

48. Caswell H (1989) Matrix population models: construction, analysis, and interpretation (Sinauer Associates Inc., Sunderland MA, USA) p 328.

